# Natural variation in fruiting body morphology in the amoeba *Dictyostelium discoideum*

**DOI:** 10.1101/2024.07.16.603515

**Authors:** Cathleen M.E. Broersma, Sean McFadyen, Elizabeth A. Ostrowski

## Abstract

Reproductive altruism, where some individuals reproduce and others do not, is considered one of the pinnacles of cooperative societies. However, the optimal level of reproductive altruism is likely to depend on inclusive fitness considerations, including the relatedness of reproducing to non-reproducing individuals, as well as the benefits and costs accruing to each, respectively. In the social amoeba *Dictyostelium discoideum*, thousands of cells aggregate to form a multicellular fruiting body. During this process, some cells die, forming a rigid stalk that supports the rest of the cells, which become viable spores. The level of stalk investment by the social group can therefore be considered a metric of altruism investment. Importantly, genetically unrelated cells can co-aggregate to produce ‘chimeric’ fruiting bodies, and selection can favour genotypes that behave selfishly by preferentially forming spores and avoiding forming the stalk. Owing to the extreme differences in fitness consequences of stalk cells versus spores, the level of altruism investment is likely to be under strong selection. Here we examined clonal fruiting body morphology in four natural populations to assess the extent to which stalk investment varies within populations and is maintained to different extents among populations. We found variation in fruiting body size and stalk investment, at both a cm-scale and between geographically isolated populations. These findings indicate the divergent evolution of altruism investment with and among populations and demonstrate widespread potential for cheating.

## Introduction

In many social groups, individuals specialise on different tasks, called division of labour. In the case of reproductive division of labour, a fraction of individuals reproduce and others aid in their reproduction (Wilson 1975; Harvell 1994). Reproductive division of labour is commonly observed in social Hymenoptera (ants, wasps, bees, and termites), where the queen reproduces, and workers (often) do not. Instead, these individuals rear the queen’s offspring, defend the nest, or forage for food (Wilson 1975; Robinson 1992; C. R. Smith et al. 2008). The specialization on different tasks is thought to enhance colony efficiency and therefore, to have allowed more complex social groups to evolve (Oster and Wilson 1978; Tsuji 1994; Reeve and Keller 1995; Nonacs and Hager 2011; Van Gestel, Vlamakis, and Kolter 2015; Cooper and West 2018).

Most multicellular organisms also show reproductive division of labour, since germline cells specialise on reproduction, and somatic cells carry out structural and functional tasks but do not themselves reproduce (Michod and Roze 2001). Although less extreme, some reproductive skew is observed in many other societies. Birds, fish, rodents, and primates (including humans) exhibit a continuum, ranging from equitable reproduction among individuals in the group (i.e., low reproductive skew) to reproduction being limited to only a few individuals (i.e., high reproductive skew) (Sherman et al. 1995; Nonacs and Hager 2011).

In clonal societies, reproductive division of labour can be readily explained by kin selection. For example, most multicellular organisms form from a single starting cell, in which case, the fitness interests of the reproductive and non-reproductive cells are aligned through very high relatedness (i.e., clonality) (Hamilton 1964; Cooper and West 2018). Here, the somatic cells’ direct fitness costs are mitigated by the indirect fitness benefits they accrue through the reproduction of clonemates. In non-clonal societies, however, reproductive division of labour can potentially be undermined by selfish cheaters, who contribute less to non-reproductive tasks and exploit cooperation by other cells (Trivers 1971; Bull and Rice 1991; Stuart A. West, Pen, and Griffin 2002; Sachs et al. 2004). Evolutionary theory therefore suggests that the level of reproductive division of labour achieved will be impacted by many factors, including relatedness, the costs and benefits associated with the behavior, and extrinsic factors (Hamilton 1964; S. A. West and Cooper 2016).

*Dictyostelium discoideum* is a soil amoeba that exhibits reproductive division of labour during its multicellular life stage. Upon starvation, thousands of individual cells aggregate to form a migratory slug and eventually a fruiting body. The fruiting body consists of approximately 80% reproductive spores and 20% non-reproductive stalk cells (Bonner and Slifkin 1949; Bonner 1982; Kessin 2001). Because the cells that form the stalk die, this act of self-sacrifice can be considered an example of altruism (Buss 1982; Armstrong 1984; DeAngelo, Kish, and Kolmes 1990). Specifically, the death of the stalk cells is thought to provide an advantage to the spores by lifting them above the soil and promoting their dispersal to better environments (J. Smith, Queller, and Strassmann 2014). The observed spore-to-stalk ratio might thus represent a balance between the indirect fitness benefits of stalk formation for related spores and the direct fitness benefits of becoming a spore oneself (Kaushik and Nanjundiah 2003). Taking into account the ecological circumstances and biophysical limitations of a fruiting body unit, there may also be an optimal ratio of cells performing both tasks where group fitness is maximized (DeAngelo, Kish, and Kolmes 1990; Matsuda and Harada 1990).

Importantly, the aggregative nature of fruiting body formation in *D. discoideum* allows genetically unrelated cells to end up in a single, ‘chimeric’ fruiting body (Buss 1982). Under these conditions, conflicts can arise over which cells become viable spores and which cells self-sacrifice to become dead stalk cells (Strassmann and Queller 2011; Strassmann, Zhu, and Queller 2000). Theory predicts that, when relatedness is sufficiently low, selection could favour cheating: individuals that preferentially form spores and avoid the costly stalk fate. Consistent with this possibility, cheating is observed in natural populations (Strassmann, Zhu, and Queller 2000) and arises readily in the laboratory (Santorelli et al. 2008).

One possibility is that the potential for cheating leads to lower levels of altruism investment (Hudson et al. 2002; Fisher, Cornwallis, and West 2013; S. A. West and Cooper 2016). Whether this does happen will depend on the mechanisms by which strains cheat, the frequency of chimerism, and the costs of stalklessness. Specifically, if cheating arises from fixed differences among strains in their spore-to-stalk ratio, called fixed allocation cheating, cheating may drive reductions in stalk height (Matapurkar and Watve 1997; Hudson et al. 2002; Kaushik and Nanjundiah 2003; Brännström and Dieckmann 2005; Cooper and West 2018; Madgwick et al. 2018). Other possibilities are that an intermediate level of altruism evolves, or coexistence of fixed allocation cheaters and strains that exhibit a high level of stalk investment (Matapurkar and Watve 1997; Brännström and Dieckmann 2005). Alternatively, if cheating results from a facultative shift to spore production, called facultative cheating, alleles that confer cheating behaviour could potentially sweep to fixation without influencing the overall level of stalk investment in the population (Ross-Gillespie et al. 2007).

Although studies have shown that *Dictyostelium* is capable of facultative cheating (Santorelli et al. 2008), little is known about whether fixed allocation cheating is common – which requires that strains vary in their intrinsic spore-stalk allocation patterns. This failure results in part from the difficulty of directly quantifying stalk allocation. Instead, researchers typically rely on indirect measurements of either stalk height or stalk volume. For example, Buttery *et al*. (2009) estimated the stalk volume based on images of upright fruiting bodies. They determined stalk length and the width at its base, midpoint, and just below the sorus and used these measurements to calculate stalk volume, assuming the geometry of a cylinder. While they did not directly report variation in stalk height among strains, they did report significant variation in the sorus-to-stalk volume ratio among seven strains sampled from a site in North Carolina, suggesting possible polymorphism in altruism investment.

Votaw and Ostrowski (2017) estimated stalk investment in populations from Texas and North Carolina in two ways. The first way involved measuring the length of the stalk in images of fruiting bodies on their sides. From these analyses, they found that strains from Texas produced larger fruiting bodies (i.e., both taller stalks and more spores) compared to strains from North Carolina, suggesting larger aggregate sizes in Texas strains. After correcting for differences in size, there was also variation among strains in their spore-to-stalk ratio, suggesting that the strains have also diverged in their level of stalk investment. To verify this finding, they generated GFP-reporters for some of the strains, in which GFP was expressed under the control of the promoter of a prespore gene, *cotB*. Using these reporter strains, they quantified the prespore and prestalk fractions of the slug (the posterior and anterior regions, respectively), and they used the ratio as an estimate of the spore-stalk allocation. In agreement with their findings from the fruiting body measurements, they again detected significant variation among strains in the spore-stalk allocation. The finding of significant variation in spore-stalk ratios among co-occurring strains is important, as it means there is an opportunity for fixed allocation cheating, where strains that commit fewer cells to the stalk can benefit from the greater stalk investment of partner strains in chimerae.

In this study, we examined fruiting body morphology across a greater range of strains and sites. In contrast to earlier studies, we examined the spore-stalk allocation of strains from the same versus different soil samples. We determined whether co-occurring strains varied in their clonal stalk investment, and we investigated whether fruiting body height and spore-to-stalk investment have evolved divergently across geographically distant sites.

These analyses address several questions. First, is fixed allocation cheating likely to be common within populations? The answer to this question requires knowing whether spore-stalk allocation variation exists at the cm-scale in the soil. Moreover, the repeated observation of spore-stalk allocation polymorphism across populations suggests that stalk-favouring and stalk-avoiding strains could co-exist. Second, how constrained is fruiting body morphology? How extreme can the relative sizes of the sorus and stalk be? While the ecological causes of any morphological divergence are not investigated, analyses of biological scaling in fruiting body dimensions can aid in identifying physical constraints on fruiting body size or proportioning that influence the evolution of altruism investment (i.e., the proportion of cells that form stalk). Third, the ability to examine intraspecific variation in altruism investment allows us to test whether these traits evolve in accordance with evolutionary theory; specifically, to test the prediction that higher altruism investment should be observed in populations where relatedness is high or cheating intensity is low (Cooper and West 2018; Madgwick et al. 2018).

## Materials and Methods

### Strain selection

The strains were collected between 2016 and 2023 by staff and students of the Ostrowski laboratory. The collection of soil and the isolation of strains are described in Kuzdzal-Fick *et al*. (2023). The strains and sites are listed in Table S1 and shown in Figure 1.

**Figure 1.**
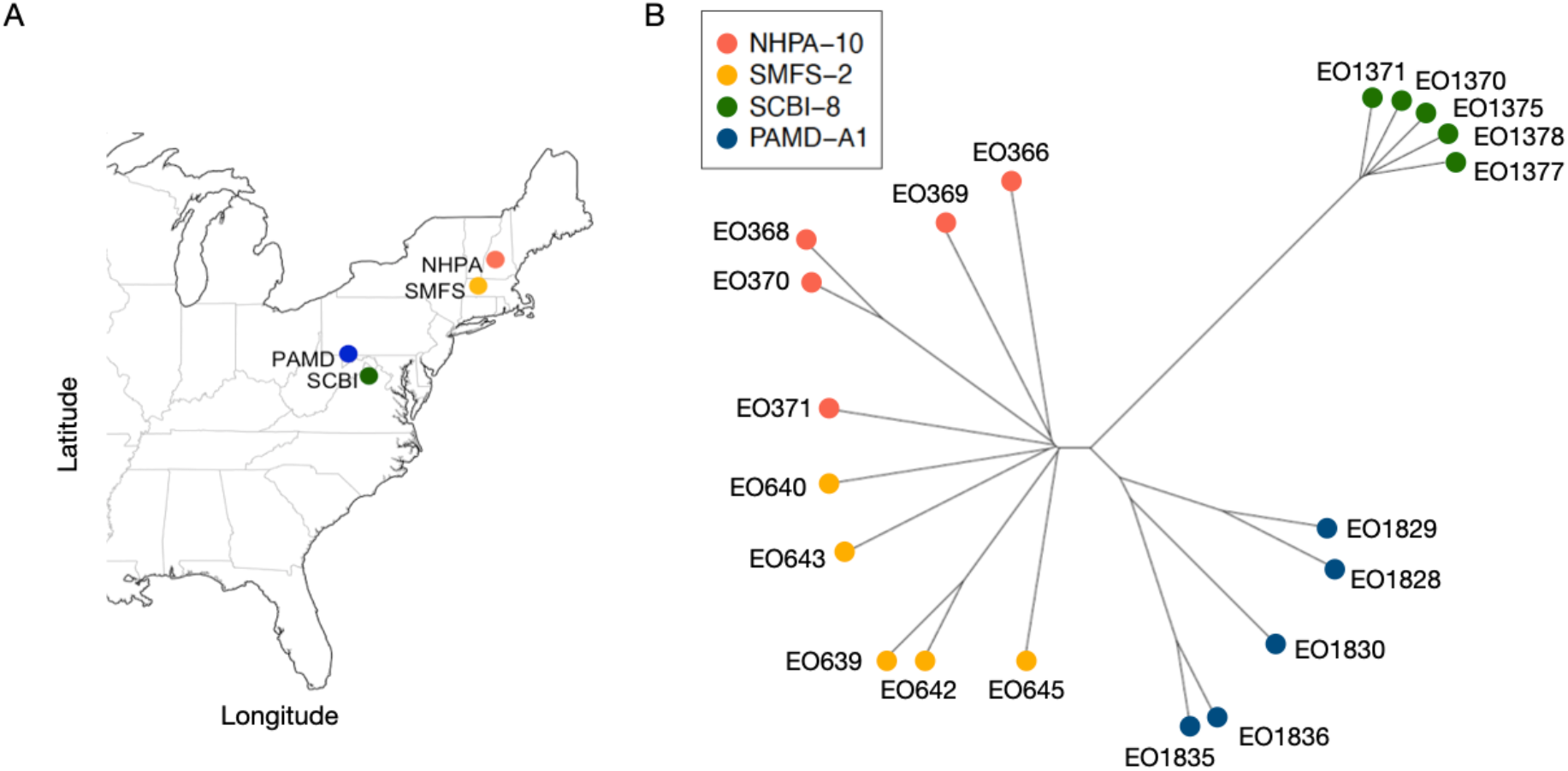
(A) Sampling locations of the four sites. (B) A distance-based tree showing genetic similarity based on genome-wide SNPs among the twenty strains.

### Strain cultivation and development

At the start of each block, we inoculated the spores from freezer stocks onto lawns of *Klebsiella pneumoniae* on SM-agar plates (SM broth, Formedium Ltd., 2% agar). After fruiting bodies had formed, we collected the spores and replated 5.0×10^5^ spores with *K. pneumoniae* on fresh SM-agar plates. We harvested the cells during mid-exponential phase and washed them three times in cold KK2 buffer (14.0 mM K_2_HPO_4_ and 3.4 mM KH_2_PO_4_, pH 6.4) via differential centrifugation at 450 x *g* for 3 min to remove the bacteria. We then resuspended the cells at a density of 1×10^8^ cells/ml in cold KK2.

For the development of the strains, we deposited a 50 μL aliquot of the cell suspension in a 3-by-3 square (∼1cm^2^) of a gridded 0.45 μM nitrocellulose filter, resulting in a total of 5.0×10^6^ cells. We transferred this filter to a 6-cm Petri dish that contained a Pall filter moistened with 1.5 mL PDF buffer (per litre: 1.5 g KCl, 1.07 g MgCl·6H_2_O, 1.8 g KH_2_PO_4_, 1.6 g K_2_HPO_4_, 0.5 g streptomycin sulphate). The Petri dishes were transferred to a plastic box lined with wet paper towels on the bottom and stored for 48 hours at 22°C in the dark. Following development, we determined the morphology of the individual fruiting bodies according to methods described in Votaw and Ostrowski (2017). Briefly, we used forceps to collect ten randomly chosen fruiting bodies. We placed each fruiting body in a separate well of a 96-well plate containing 100 μL of spore detergent (0.1 % IGEPAL in KK2 buffer with 10 mM EDTA) and imaged the wells at 50x magnification. From the images, we measured the length of the stalks in ImageJ. Since most stalks were bent, we used the “segmented line” function to measure the length of the stalk from the bottom of the spore head to the top of the basal disk. After imaging, we used a pipette to disperse the spores and lyse any remaining cells. We used an automated cell counter (Cell Countess II, Thermo Fisher) to calculate the number of spores and the average spore size in each fruiting body. In total, we measured the stalk length, the number of spores produced, and the average spore size in 570 fruiting bodies (19 strains x 3 blocks x 10 fruiting bodies).

### Statistical analyses

We performed all analyses in R version 4.2.1 (R Core Team 2023). We excluded strain EO640 from the analyses as it showed aberrant fruiting body formation, with most aggregates not progressing beyond the slug stage within 48 hours.

We tested for variation in stalk height, spore number and spore size in separate generalized linear mixed models (GLMMs). Each model was fit using maximum likelihood with the ‘glmmTMB’ function. We included site as a fixed effect and assessed its significance through a type II Wald chi-square test. The models that examined stalk height and spore number had a Gaussian error structure, and the model that examined average spore size had a Gamma error structure. We included strain ID and block as random effects and assessed their significance through single deletions of terms, comparing the reduced models and full model using a likelihood ratio test.

We examined the relationship between stalk height and spore number using the ‘sma’ function from the ‘smatr’ package (Warton et al. 2012). This regression method allows one to examine a scaling relationship between two variables where neither of them is predicting the other. We tested for variation in the slope using the formula spore number ∼ stalk height * group. We tested for variation in intercept using the formula spore number ∼ stalk height + group which assumes common slopes. We tested for variation in the shift along the common axis using the formula spore number ∼ stalk height + group, type = “shift”. In each analysis, ‘group’ was either site or strain ID when we tested for variation among sites or among strains within a site respectively. We performed a Sidak correction for multiple comparisons and set the parameter ‘robust=TRUE’ to fit the line using Huber’s M estimation to downweight outliers (Warton et al. 2012). We assessed the significance of the slope and intercept using the likelihood ratio statistic from the ‘summary’ function. We assessed the significance of the position along the common axis using the Wald statistics from the ‘summary’ function. We carried out the principal components analysis using the ‘prcomp’ function from the ‘stats’ package with centering and scaling.

### Relationships between spore inequity (as a metric of cheating) and genetic distance, and stalk investment

To test the prediction that higher altruism investment is found in populations where relatedness is high or cheating intensity is low, we examined the relationship between the average spore-to-stalk ratio (quantified in this study) and the average genetic distance or magnitude of cheating found within a site (quantified in Broersma and Ostrowski, in prep.) respectively. Specifically, in a prior study, we quantified spore inequity among pairs of strains in chimerae as a metric of cheating, using a subset of the strain pairs used here (*N*=6-7 strain pairs tested per site) (Broersma and Ostrowski, in prep.). We quantified the magnitude of spore inequity as the deviation from the ratio of the two strains in the spores compared to that of the cells before development—which was 0.5 because of equal mixing of strains. For example, a spore inequity value of 0.1 indicates that the ratio of strains shifted from 50-50 in the cells to 60-40 in the spores. We tested the significance of the relationship between spore inequity magnitude and spore-to-stalk ratio within sites using a one-tailed Pearson’s correlation test. We used the same test to examine the relationship between the average genetic distance and the spore-to-stalk ratio within sites.

## Results

### Average fruiting body dimensions

We measured three traits of fruiting body morphology: stalk height, number of spores, and average spore size. Across all strains, the average fruiting body had a stalk height of 3.39±0.07 mm, contained a total of 3.69×10^4^±2.4×10^3^ spores, and the spores had an average size of 1.60±0.03 mm (grand mean across *N*=19 strains). On each filter, we deposited 5×10^6^ cells and collected an average of 4.27×10^6^ spores following development, which indicates that ∼85% of cells became spores.

### Within- and between-site variation in fruiting body morphology

There was significant variation in stalk height and spore size, but not spore number, across sites (Figure 2; Table S2). In addition to differences across sites, there was significant variation among strains within sites for all three traits (Figure 2; Table S2). Together, these observations demonstrate both polymorphisms within sites and divergence across sites in fruiting body morphology.

**Figure 2.**
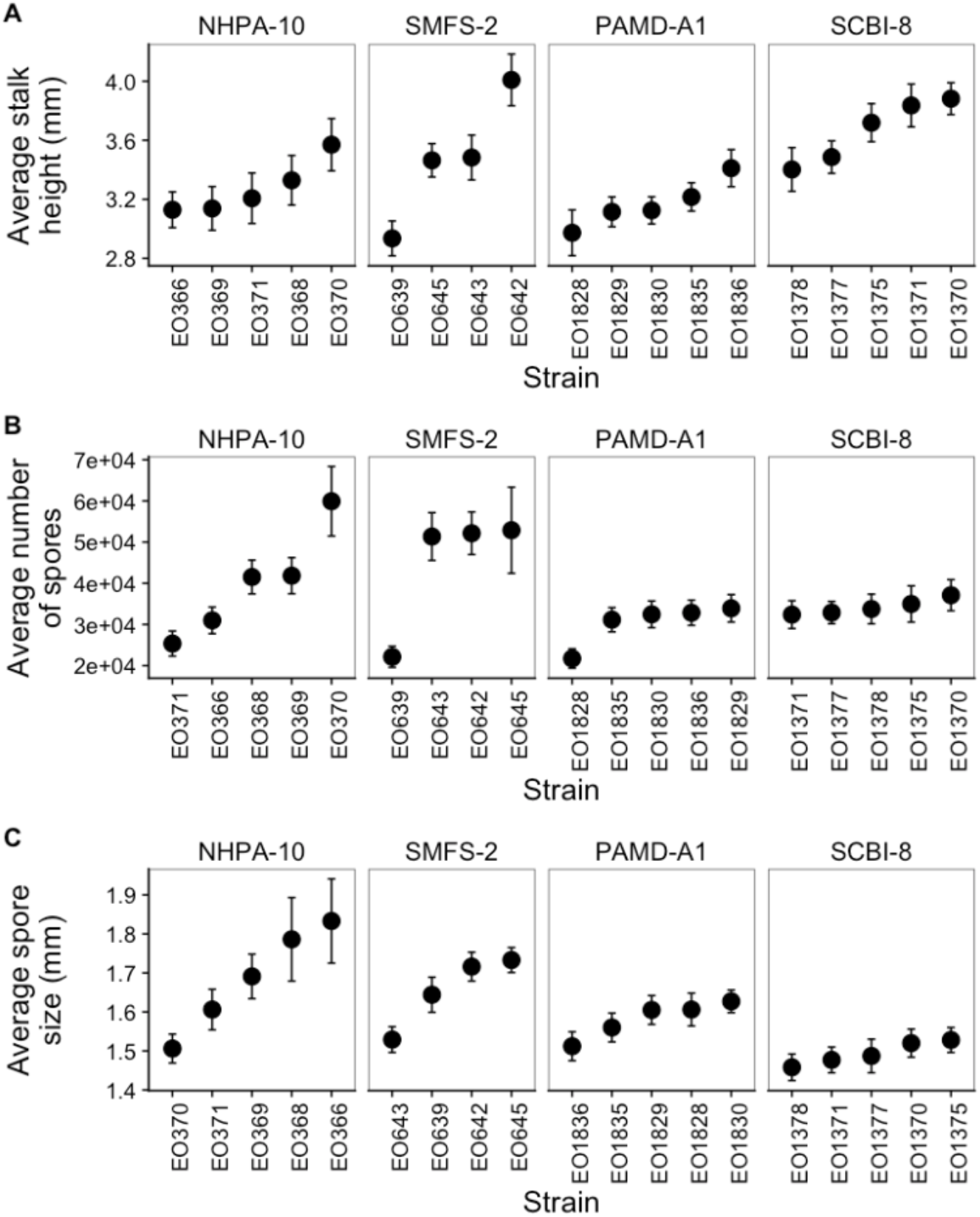
Fruiting body traits (A) stalk height, (B) spore number, and (C) average spore size for four to five strains from four sites. Each point represents the grand mean for a single strain based on 30 fruiting bodies (=3 blocks x 10 fruiting bodies). The error bars represent one standard error of the mean (based on *N=*3 blocks).

### Analysis of the biological scaling relationship between spore number and stalk height

We examined the biological scaling relationship between spore number and stalk height to get an estimate of the level of stalk investment and test if this estimate varies among and within sites. This method allowed us to disentangle absolute changes in size from proportional changes that can reflect differences in spore-to-stalk investment (Warton et al. 2012). For example, one strain might produce fruiting bodies that contain more spores *and* taller stalks than another strain; this strain might simply produce larger fruiting bodies, without a change in the relative allocation of the spores versus the stalk. This distinction is relevant because we are interested in whether strains vary in their altruism investment, that is, the fraction of cells allocated to the reproductive versus non-reproductive cell fate. Figure 3A illustrates potential differences in the relationship between spore number and stalk height in four hypothetical strains.

**Figure 3.**
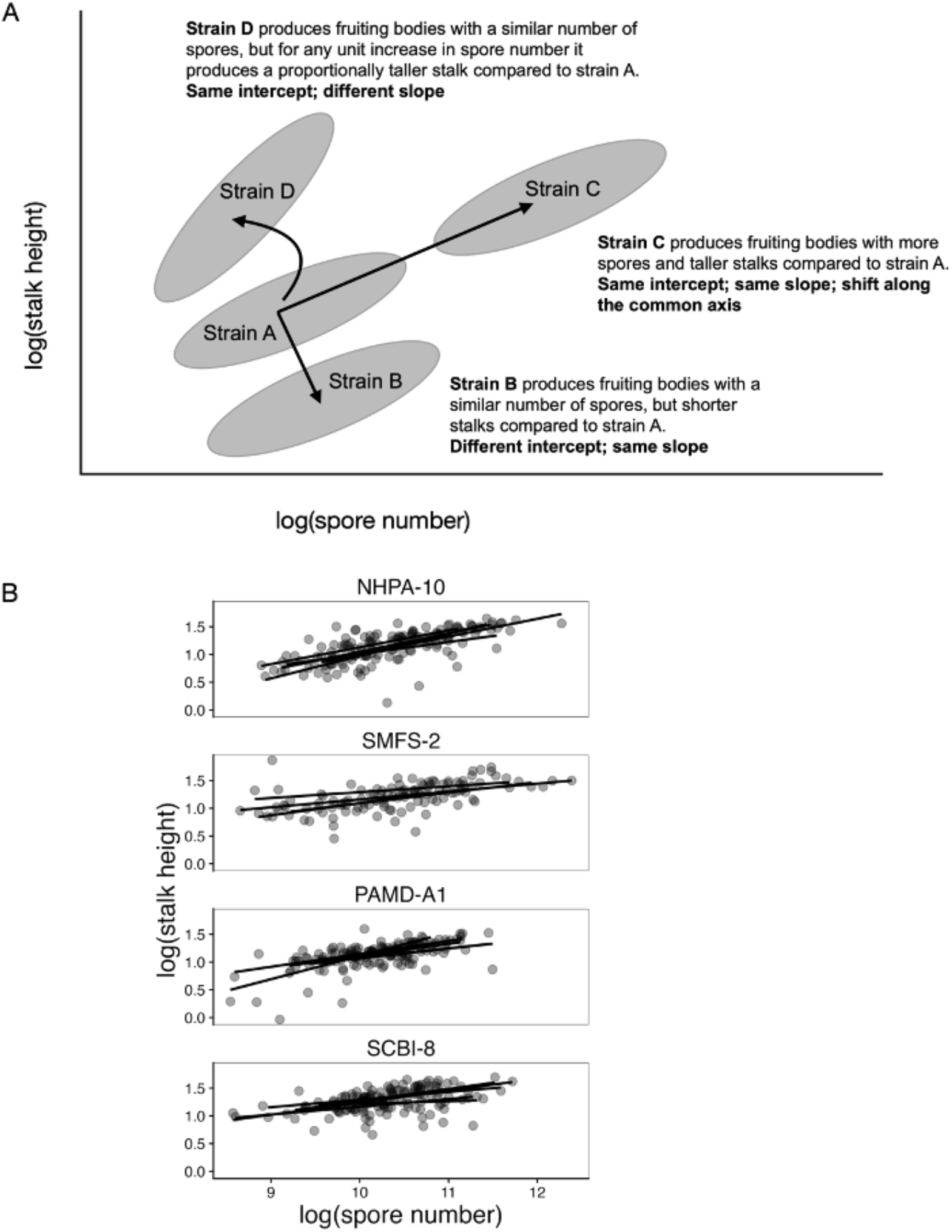
(A) The relationship between spore number and stalk height in four hypothetical strains demonstrates how to distinguish between differences in stalk (altruism) investment and fruiting body size. (B) The observed relationship between spore number and stalk height in strains from four sites. Each point indicates a single fruiting body. Lines indicate the best fit (based on *N*=30; 10 fruiting bodies x 3 blocks).

### Within- and between-site variation in stalk investment and group size

Apart from strains EO642 and EO1377, all strains showed a significant positive relationship between spore number and stalk height (Figure 3B and Table S3). Moreover, we found significant variation in the degree to which stalk investment changes with size (i.e., differences in slope), stalk investment (i.e., differences in intercept), and group size (i.e., differences in the position along the common axis) both within and across sites (Table S4). Specifically, NHPA-10 and PAMD-A1 showed significant variation in spore-stalk allocation among strains. Strains from SMFS-2 showed variation in size but not spore-stalk allocation. Notably, strains from SCBI-8 showed no significant variation in any of the three traits, meaning they were morphologically similar (Table S4).

### Principal components analysis

To quantify the relative contributions of stalk investment versus fruiting body size to variation in fruiting body morphology, we also performed a principal component analysis (PCA). PC1 was influenced most strongly by stalk height and spore number, both in a positive direction, and explained 56.0% of the variance (Table 1). As expected, this suggests that PC1 predominantly captures the variation in size. Interestingly, spore size was negatively related to spore number and stalk height (albeit more weakly), suggesting that spore size might trade off with spore number. Because PC1 mostly captures the variation in size, this allowed us to examine the other PCs for variation in stalk investment after we have corrected for variation in size in PC1. PC2 was influenced most strongly by spore size and explained 31.0% of the variance. PC3 was influenced most strongly by stalk height and spore number, but in opposite directions, and explained 13.0% of the variance. Thus, PC1 primarily captured the variation in size, PC2 primarily captured the variation in average spore size, and PC3 captured the variation in spore number versus stalk size *after controlling for size*. Therefore, high values of PC1 represent fruiting bodies of greater size (i.e., more spores *and* taller stalks), whereas high values of PC3 represent fruiting bodies with reduced stalk investment (i.e., more spores but shorter stalks).

**Table 1.** Factor loadings of the principal component analysis.

| Trait | PC1 | PC2 | PC3 |
| --- | --- | --- | --- |
| Spore number | 0.647 | -0.359 | 0.672 |
| Stalk height | 0.683 | -0.118 | -0.721 |
| Average spore size | -0.338 | -0.926 | -0.169 |
| % Variance | 56.0 | 31.0 | 13.0 |

### Is greater cheating and lower relatedness associated with decreases in stalk investment?

Populations are likely to vary in their cheating load, reflecting their different ecological and genetic circumstances (Bruce et al. 2017; Butaitė, Kramer, and Kümmerli 2021). For example, populations where relatedness is high or low should permit cheating to different extents (low and high, respectively). Where cheating is pervasive, clonal stalk investment may be lower, reflecting a different evolutionary balance between the benefits of stalk production and the costs of cheating (Hudson et al. 2002). Put another way, we expect that populations with high relatedness and/or low intensity of cheating might maintain higher stalk investment. To see whether this is true, we examined the relationship between average relatedness, cheating intensity, and the spore-to-stalk ratio. For cheating intensity, we used the estimates of mean spore inequity from a prior study (Broersma and Ostrowski in prep.) and we used the above-obtained estimate of PC3 as a metric for the spore-to-stalk ratio. We found that the relationship between spore inequity and the spore-to-stalk ratio was not significant (Figure 4A; one-tailed Pearson’s correlation *r*=-0.59, df=2, *P*=0.79), but the correlation was moderate in magnitude and negative, as predicted. Similarly, the relationship between genetic distance and the spore-to-stalk ratio was not significant (Figure 4B; one-tailed Pearson’s correlation *r*=-0.87, df=2, *P*=0.07), but, again, the correlation was strong in magnitude and negative, as predicted.

**Figure 4.**
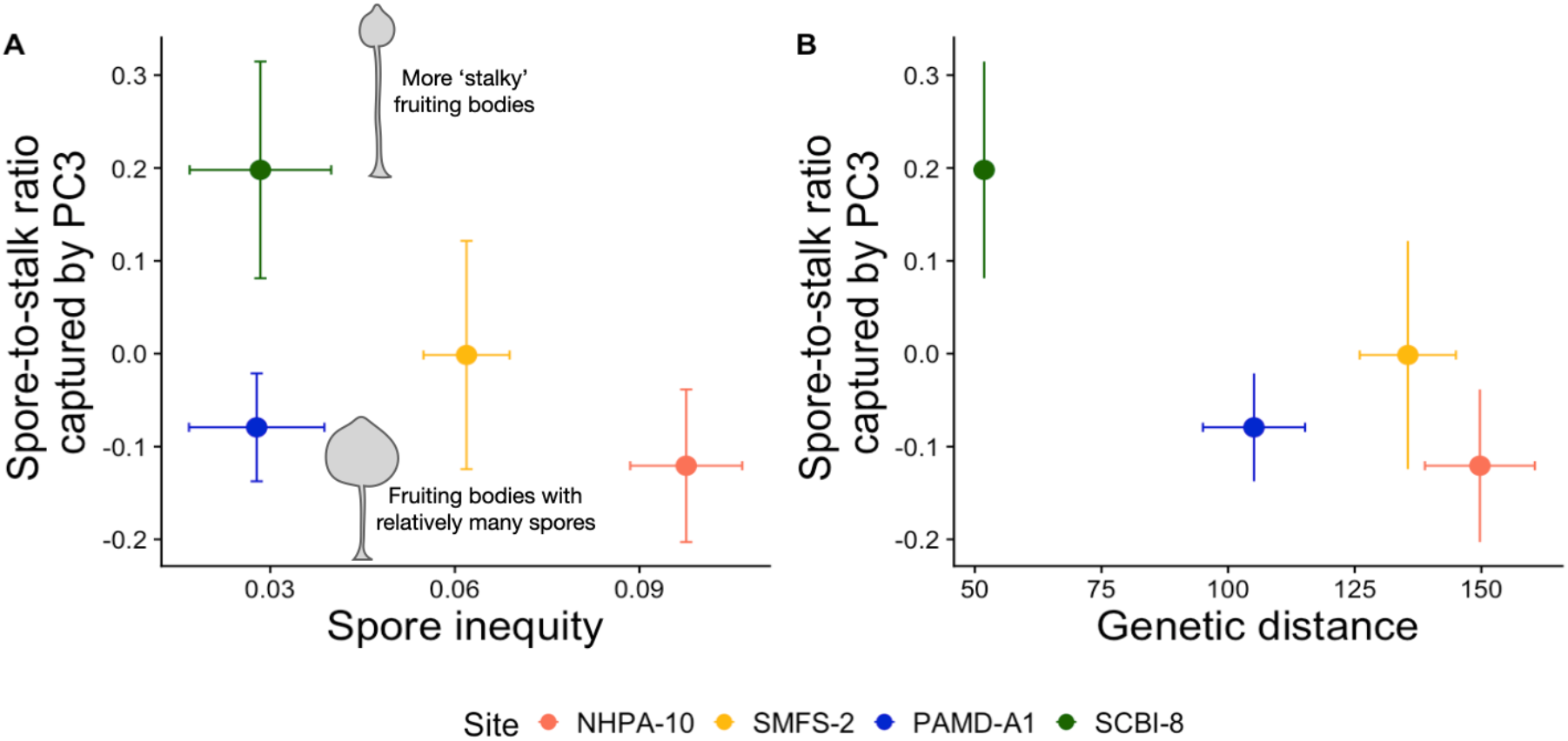
(A) Relationship between mean spore inequity and clonal spore-to-stalk ratio for four populations. PC3 was used as a proxy for greater spore investment (see text). High values of PC3 indicate fruiting bodies with relatively tall stalks after controlling for overall size. The average magnitude of spore inequity within a site was quantified in another study (Broersma and Ostrowski, in prep.; see the Methods section for how spore inequity was measured). (B) The relationship between genetic distance and spore-to-stalk ratio for four populations. In (A) and (B) the error bars represent one standard error of the mean.

## Discussion

Species that live in social groups and exhibit reproductive division of labour show a continuum in the level of altruism investment, i.e., the fraction of individuals that forgo reproduction themselves to promote that of others in their group. Theory and empirical work suggest that the level of altruism investment will be influenced by many factors, including the relatedness between the non-reproductive and reproductive individuals, as well as the costs and benefits associated with either subgroup (Arnold, Owens, and Goldizen 2005; Fisher, Cornwallis, and West 2013; Ferguson-Gow et al. 2014; S. A. West and Cooper 2016; Schiessl et al. 2019). Here we measured spore-to-stalk allocation as a proxy for altruism investment in four populations of the social amoeba *Dictyostelium discoideum*. We found variation in spore-to-stalk allocation among geographically isolated populations as well as among strains isolated from a 10-by-10 cm plot of soil. These results suggest both population divergence in the level of altruism investment and polymorphism in traits important for social interactions.

There was a consistently positive correlation between stalk height and spore number, meaning that variation in stalk height was in part attributable to variation in multicellular size. One explanation for increased fruiting body size could be the need to produce taller stalks in some environments, as discussed by Votaw and Ostrowski (2017). Larger fruiting bodies will lift spores further off the ground, without requiring any greater per capita investment in stalk production. In other words, selection for taller stalks may indirectly select for larger size. Future work would benefit from examining fruiting body size across environments in which taller or shorter stalks could be favoured, for example, in more or less humid environments (Bonner and Shaw 1957).

Two of the four sites showed significant variation in stalk investment among strains within a site. Together with two prior studies that examined the same set of traits, these findings suggest that stalk-avoiding and stalk-favouring strains may often coexist within a population (Buttery et al. 2009; Votaw and Ostrowski 2017). This possibility is consistent with theoretical work showing that under certain conditions, cheaters and cooperators can coexist in a stably oscillating population (Matapurkar and Watve 1997; Brännström and Dieckmann 2005). In addition, genomic analyses revealed signatures of negative frequency-dependent selection in candidate genes that mediate cheating behaviours, suggesting that multiple alleles are maintained in populations (Ostrowski et al. 2015).

Fixed allocation differences among strains were shown previously to contribute to inequities in spore production when two strains form a chimera (Buttery et al. 2009; Broersma and Ostrowski, in prep.) However, so far, no studies have determined whether this variation exists at the centimetre scale—i.e., the spatial scales over which strains are likely to interact in the soil. This study is unique in examining social trait diversity over these small, biologically relevant spatial scales, and it greatly expands the number of populations in which this variation has been demonstrated, revealing substantial variation on which selection can act.

Importantly, the level and variation in altruism investment in this system will likely depend on the frequency of chimeric fruiting body formation in nature and the relationship between stalk height and fitness (dispersal success). Unfortunately, both factors are not known with certainty. However, one study showed that 63% of small soil samples – each just 6 mm in diameter – harboured more than one genotype of *D. discoideum* (Fortunato et al. 2003), suggesting that genetically different strains are likely to encounter one another frequently in nature. With respect to the importance of stalk height to dispersal success, one study showed that spores from intact fruiting bodies are better dispersed by flies compared to spores from knocked-over (‘stalkless’) fruiting bodies (J. Smith, Queller, and Strassmann 2014). However, no studies have determined whether existing variation in stalk height impacts dispersal success. Though this will likely be challenging to test, future work would benefit from better estimates of the costs associated with lower stalk investment, given that natural variation in this trait is observed within and across populations, as well as in response to chimeric development (Votaw and Ostrowski 2017; Kuzdzal-Fick et al. 2023).

Assessing the extent to which social traits vary across and within natural populations is important for understanding how these traits evolve in nature. In this study, we have not addressed what evolutionary forces promote and maintain intraspecific diversity. However, we have shown that more and less altruistic phenotypes coexist at a cm-scale in nature and that geographically distant populations have diverged in their levels of altruism investment. These factors can impact the nature and outcome of social interactions between locally co-occurring genotypes and genotypes that were geographically isolated but re-introduced.

## Acknowledgements

The authors would like to thank Smith MacLeish Field Station, Proctor Academy, Forbes State Forest, and Smithsonian Conservation Biology Institute for permission to collect soil samples. We thank Michael Miller, Maria Polo Prieto, Scott Clark, and Dave Neely for assistance with soil collection and strain isolation. This work was funded by grants to EAO from the US National Science Foundation DEB-1557023 and the Marsden Fund, administered by Te Apārangi Royal Society of New Zealand, as well as a Massey University Doctoral Scholarship to CMB.

## Author contributions

CMB and EAO conceived the idea and designed the experiments. CMB and SM collected the data. CMB analysed the data. CMB and EAO wrote the manuscript. CMB, EAO and SM approved the final version.

## Data accessibility

Data and scripts are available upon request.

## Literature cited

Armstrong, Doug P. 1984. “Why Don’t Cellular Slime Molds Cheat?” Journal of Theoretical Biology. 10.1016/s0022-5193(84)80006-5.

Arnold, Kathryn E., Ian P. F. Owens, and Anne W. Goldizen. 2005. “Division of Labour within Cooperatively Breeding Groups.” Behaviour 142 (11/12): 1577–90.

Bonner, J. T. 1982. “Evolutionary Strategies and Developmental Constraints in the Cellular Slime Molds.” The American Naturalist 119 (4): 530–52.

Bonner, J. T., and M. J. Shaw. 1957. “The Role of Humidity in the Differentiation of the Celular Slime Molds.” Journal of Cellular and Comparative Physiology 50 (1): 145–53.

Bonner, J. T., and M. K. Slifkin. 1949. “A Study of the Control of Differentiation: The Proportions of Stalk and Spore Cells in the Slime Mold Dictyostelium Discoideum.” American Journal of Botany 36 (10): 727–34.

Brännström, A., and U. Dieckmann. 2005. “Evolutionary Dynamics of Altruism and Cheating among Social Amoebas.” Proceedings. Biological Sciences / The Royal Society 272 (1572): 1609–16.

Bruce, John B., Guy A. Cooper, Hélène Chabas, Stuart A. West, and Ashleigh S. Griffin. 2017. “Cheating and Resistance to Cheating in Natural Populations of the Bacterium Pseudomonas Fluorescens.” Evolution; International Journal of Organic Evolution 71 (10): 2484–95.

Bull, J. J., and W. R. Rice. 1991. “Distinguishing Mechanisms for the Evolution of Co-Operation.” Journal of Theoretical Biology 149 (1): 63–74.

Buss, L. W. 1982. “Somatic Cell Parasitism and the Evolution of Somatic Tissue Compatibility.” Proc. Natl. Acad. Sci. U S A 79: 5337–41.

Butaitė, Elena, Jos Kramer, and Rolf Kümmerli. 2021. “Local Adaptation, Geographical Distance and Phylogenetic Relatedness: Assessing the Drivers of Siderophore-Mediated Social Interactions in Natural Bacterial Communities.” Journal of Evolutionary Biology 34 (8): 1266–78.

Buttery, N. J., D. E. Rozen, J. B. Wolf, and C. R. L. Thompson. 2009. “Quantification of Social Behavior in D. Discoideum Reveals Complex Fixed and Facultative Strategies.” Curr. Biol. 19 (16): 1373–77.

Cooper, G. A., and S. A. West. 2018. “Division of Labour and the Evolution of Extreme Specialization.” Nature Ecology & Evolution 2 (7): 1161–67.

DeAngelo, M. J., V. M. Kish, and S. A. Kolmes. 1990. “Altruism, Selfishness, and Heterocytosis in Cellular Slime Molds.” Ethology Ecology & Evolution 2 (4): 439–43.

Ferguson-Gow, Henry, Seirian Sumner, Andrew F. G. Bourke, and Kate E. Jones. 2014. “Colony Size Predicts Division of Labour in Attine Ants.” Proceedings. Biological Sciences / The Royal Society 281 (1793). 10.1098/rspb.2014.1411.

Fisher, Roberta M., Charlie K. Cornwallis, and Stuart A. West. 2013. “Group Formation, Relatedness, and the Evolution of Multicellularity.” Current Biology: CB 23 (12): 1120–25.

Fortunato, A., J. E. Strassmann, L. Santorelli, and D. C. Queller. 2003. “Co-Occurrence in Nature of Different Clones of the Social Amoeba, Dictyostelium Discoideum.” Molecular Ecology 12 (4): 1031–38.

Hamilton, W. D. 1964. “The Genetical Evolution of Social Behaviour. I.” Journal of Theoretical Biology 7 (1): 1–16.

Harvell, C. Drew. 1994. “The Evolution of Polymorphism in Colonial Invertebrates and Social Insects.” The Quarterly Review of Biology 69 (2): 155–85.

Hudson, Aukema, Rispe, and Roze. 2002. “Altruism, Cheating, and Anticheater Adaptations in Cellular Slime Molds.” The American Naturalist 160 (1): 31.

Kaushik, Sonia, and Vidyanand Nanjundiah. 2003. “Evolutionary Questions Raised by Cellular Slime Mould Development.” Proc Indian Natl Sci Acad B69: 825–52.

Kessin, R. H. 2001. Dictyostelium. Evolution, Cell Biology, and the Development of Multicellularity. Edited by J. B. L. Bard, P. W. Barlow, and D. L. Kirk. Developmental and Cell Biology. Cambridge University Press.

Kuzdzal-Fick, J. J., A. Moreno, C. M. E. Broersma, T. F. Cooper, and E. A. Ostrowski. 2023. “From Individual Behaviors to Collective Outcomes: Fruiting Body Formation in Dictyostelium as a Group-Level Phenotype.” Evolution; International Journal of Organic Evolution 77 (3): 731–45.

Madgwick, P. G., B. Stewart, L. J. Belcher, C. R. L. Thompson, and J. B. Wolf. 2018. “Strategic Investment Explains Patterns of Cooperation and Cheating in a Microbe.” Proceedings of the National Academy of Sciences of the United States of America 115 (21): E4823–32.

Matapurkar, Anagha K., and Milind G. Watve. 1997. “Altruist Cheater Dynamics in Dictyostelium: Aggregated Distribution Gives Stable Oscillations.” The American Naturalist 150 (6): 790–97.

Matsuda, H., and Y. Harada. 1990. “Evolutionarily Stable Stalk to Spore Ratio in Cellular Slime Molds and the Law of Equalization in Net Incomes.” Journal of Theoretical Biology 147 (3): 329–44.

Michod, Richard E., and Denis Roze. 2001. “Cooperation and Conflict in the Evolution of Multicellularity.” Heredity 86: 1–7.

Nonacs, P., and R. Hager. 2011. “The Past, Present and Future of Reproductive Skew Theory and Experiments.” Biological Reviews of the Cambridge Philosophical Society 86 (2): 271–98.

Oster, G. F., and E. O. Wilson. 1978. Caste and Ecology in the Social Insects. Princeton University Press.

R Core Team. 2023. R: A Language and Environment for Statistical Computing (version 2023.03.1). https://www.R-project.org/.

Reeve, H. K., and L. Keller. 1995. “Partitioning of Reproduction in Mother-Daughter versus Sibling Associations: A Test of Optimal Skew Theory.” The American Naturalist 145 (1): 119–32.

Robinson, G. E. 1992. “Regulation of Division of Labor in Insect Societies.” Annual Review of Entomology 37: 637–65.

Ross-Gillespie, Adin, Andy Gardner, Stuart A. West, and Ashleigh S. Griffin. 2007. “Frequency Dependence and Cooperation: Theory and a Test with Bacteria.” The American Naturalist 170 (3): 331–42.

Sachs, J. L., U. G. Mueller, T. P. Wilcox, and J. J. Bull. 2004. “The Evolution of Cooperation.” The Quarterly Review of Biology 79 (2): 135–1160.

Santorelli, Lorenzo A., Christopher R. L. Thompson, Elizabeth Villegas, Jessica Svetz, Christopher Dinh, Anup Parikh, Richard Sucgang, et al. 2008. “Facultative Cheater Mutants Reveal the Genetic Complexity of Cooperation in Social Amoebae.” Nature 451 (7182): 1107–10.

Schiessl, Konstanze T., Adin Ross-Gillespie, Daniel M. Cornforth, Michael Weigert, Colette Bigosch, Sam P. Brown, Martin Ackermann, and Rolf Kümmerli. 2019. “Individual- versus Group-Optimality in the Production of Secreted Bacterial Compounds.” Evolution; International Journal of Organic Evolution 73 (4): 675–88.

Sherman, P. W., E. A. Lacey, H. K. Reeve, and L. Keller. 1995. “The Eusociality Continuum.” Behavioral Ecology 6: 102–8.

Smith, C. R., A. L. Toth, A. V. Suarez, and G. E. Robinson. 2008. “Genetic and Genomic Analyses of the Division of Labour in Insect Societies.” Nature Reviews. Genetics 9 (10): 735–48.

Smith, J., D. C. Queller, and J. E. Strassmann. 2014. “Fruiting Bodies of the Social Amoeba Dictyostelium Discoideum Increase Spore Transport by Drosophila.” BMC Evolutionary Biology 14 (May): 105.

Strassmann, J. E., and D. C. Queller. 2011. “How Social Evolution Theory Impacts Our Understanding of Development in the Social Amoeba Dictyostelium.” Development, Growth & Differentiation 53 (4): 597–607.

Strassmann, J. E., Y. Zhu, and D. C. Queller. 2000. “Altruism and Social Cheating in the Social Amoeba Dictyostelium Discoideum.” Nature 408 (6815): 965–67.

Trivers, Robert L. 1971. “The Evolution of Reciprocal Altruism.” The Quarterly Review of Biology 46 (1): 35– 57.

Tsuji, Kazuki. 1994. “Inter-Colonial Selection for the Maintenance of Cooperative Breeding in the Ant Pristomyrmex Pungens: A Laboratory Experiment.” Behavioral Ecology and Sociobiology 35 (2): 109–13.

Van Gestel, Jordi, Hera Vlamakis, and Roberto Kolter. 2015. “Division of Labor in Biofilms: The Ecology of Cell Differentiation.” In Microbial Biofilms, 67–97. Washington, DC, USA: ASM Press.

Votaw, H. R., and E. A. Ostrowski. 2017. “Stalk Size and Altruism Investment within and among Populations of the Social Amoeba.” Journal of Evolutionary Biology 30 (11): 2017–30.

Warton, D. I., R. A. Duursma, D. S. Falster, and S. Taskinen. 2012. “Smatr 3 - an R Package for Estimation and Inference about Allometric Lines.” Methods in Ecology and Evolution / British Ecological Society 3 (2): 257– 59.

West, S. A., and G. A. Cooper. 2016. “Division of Labour in Microorganisms: An Evolutionary Perspective.” Nature Reviews. Microbiology 14 (11): 716–23.

West, Stuart A., Ido Pen, and Ashleigh S. Griffin. 2002. “Cooperation and Competition between Relatives.” Science 296 (5565): 72–75.

Wilson, E. O., ed. 1975. Sociobiology: The New Synthesis. The Belknap Press of Harvard University Press Cambridge, Massachusetts, and London, England.

