## Supplementary material for "Natural variation in fruiting body morphology in the amoeba *Dictyostelium discoideum*"

Table S1. List of strains and their sampling sites and GPS coordinates.

| Strain ID | Origin | Location ID | GPS Coordinates |
| --- | --- | --- | --- |
| EO366, EO368, EO369, EO370, EO371 | New Hampshire Proctor Academy, NH | NHPA-10 | 43°45’207, -71°82’578 |
| EO639, EO640, EO642, EO643, EO645 | Smith College MacLeish Field Station, MA | SMFS-2 | 42°44’914, -72°68’204 |
| EO1370, EO1371, EO1375, EO1377, EO1378 | Smithsonian Conservation Biology Institute, VA | SCBI-8 | 38°89’344, -78°14’647 |
| EO1828, EO1829, EO1830, EO1835, EO1836 | Pennsylvania Mount Davis, PA | PAMD-A1 | 39°78’546, -79°17’421 |

**Table S2**. Summary of the results of the mixed models that examined variation in fruiting body morphology within and across sites.

| Effects | Stalk height | | | Spore number | | | | Spore size | | |
| --- | --- | --- | --- | --- | --- | --- | --- | --- | --- | --- |
| Fixed effects | df | $\chi^{2}$ | P($\chi^{2}$) | df | $\chi^{2}$ | P($\chi^{2}$) | df | | $\chi^{2}$ | P($\chi^{2}$) |
| Site ID | 3 | 13.126 | 0.004 | 3 | 2.678 | 0.44 | 3 | | 18.827 | <0.001 |
| Random effects | df | $\chi^{2}$ | P($\chi^{2}$) | df | $\chi^{2}$ | P($\chi^{2}$) | df | | $\chi^{2}$ | P($\chi^{2}$) |
| Strain ID | 1 | 23.095 | <0.001 | 1 | 33.8 | <0.001 | 1 | | 9.546 | 0.002 |
| Block | 1 | 113.71 | <0.001 | 1 | 6.523 | 0.011 | 1 | | 48.88 | <0.001 |

Table S3. The slope, intercept, and correlation coefficient (*r*) of the relationship between spore number and stalk height.

| Strain ID | Site | Slope | Intercept | r | P |
| --- | --- | --- | --- | --- | --- |
| EO1370 | SCBI-8 | 0.294 | -2.503 | 0.49 | 0.006 |
| EO1371 | SCBI-8 | 0.436 | -2.483 | 0.51 | 0.004 |
| EO1375 | SCBI-8 | 0.354 | -2.513 | 0.74 | <0.001 |
| EO1377 | SCBI-8 | 0.395 | -2.592 | 0.10 | 0.60 |
| EO1378 | SCBI-8 | 0.403 | -2.602 | 0.39 | 0.035 |
| EO1828 | PAMD-A1 | 0.329 | -2.607 | 0.67 | <0.001 |
| EO1829 | PAMD-A1 | 0.386 | -2.703 | 0.49 | 0.006 |
| EO1830 | PAMD-A1 | 0.337 | -2.687 | 0.46 | 0.013 |
| EO1835 | PAMD-A1 | 0.284 | -2.632 | 0.72 | <0.001 |
| EO1836 | PAMD-A1 | 0.382 | -2.597 | 0.67 | <0.001 |
| EO366 | NHPA-10 | 0.430 | -2.668 | 0.79 | <0.001 |
| EO368 | NHPA-10 | 0.494 | -2.721 | 0.82 | <0.001 |
| EO369 | NHPA-10 | 0.418 | -2.774 | 0.49 | 0.006 |
| EO370 | NHPA-10 | 0.385 | -2.743 | 0.85 | <0.001 |
| EO371 | NHPA-10 | 0.456 | -2.576 | 0.56 | 0.001 |
| EO639 | SMFS-2 | 0.314 | -2.588 | 0.61 | <0.001 |
| EO642 | SMFS-2 | 0.403 | -2.597 | 0.24 | 0.10 |
| EO643 | SMFS-2 | 0.403 | -2.738 | 0.47 | 0.009 |
| EO645 | SMFS-2 | 0.200 | -2.658 | 0.70 | <0.001 |

Table S4. A summary of the models that tested if sites (top table) and strains within a site (bottom table) showed significant variation in the slope, intercept, and position along the common axis, of the relationship between spore number and stalk height.

| Term | df | $\boldsymbol{\chi}^{\mathbf{2}}$or F | P($\boldsymbol{\chi}^{\mathbf{2}}$ or F) |
| --- | --- | --- | --- |
| Slope | 3 | 9.593 | 0.022 |
| Intercept | 3 | 55.95 | <0.001 |
| Position along the common axis | 3 | 21.36 | <0.001 |

| Term | Site | df | $\boldsymbol{\chi}^{\mathbf{2}}$ or F | P($\boldsymbol{\chi}^{\mathbf{2}}$ or F) |
| --- | --- | --- | --- | --- |
| Slope  (=the degree to which stalk | NHPA-10 | 4 | 3.29 | 0.51 |
|  | SMFS-2 | 3 | 14.00 | 0.003 |
| investment changes with size) | PAMD-A1 | 4 | 4.10 | 0.40 |
|  | SCBI-8 | 4 | 3.65 | 0.46 |
| Intercept | NHPA-10 | 4 | 25.14 | <0.001 |
| (=stalk investment) | SMFS-2 | 3 | 4.39 | 0.22 |
|  | PAMD-A1 | 4 | 14.48 | 0.006 |
|  | SCBI-8 | 4 | 8.07 | 0.09 |
| Shift along the common axis | NHPA-10 | 4 | 8.28 | 0.08 |
| (=size) | SMFS-2 | 3 | 26.69 | <0.001 |
|  | PAMD-A1 | 4 | 5.07 | 0.28 |
|  | SCBI-8 | 4 | 4.71 | 0.32 |
